# MethylSeqLogo: DNA methylation smart sequence logos

**DOI:** 10.1101/2022.11.05.515271

**Authors:** Fei-Man Hsu, Paul Horton

## Abstract

**Background:** Sequence logos can effectively visualize position specific base preferences evident in a collection of binding sites of some transcription factor. But those preferences usually fall far short of fully explaining binding specificity. Interestingly, some transcription factors bind sites of potentially methylated DNA. For example, MYC binds `CpG` sites. This may increase binding specificity as such sites are 1) highly under-represented in the genome, and 2) offer additional, tissue specific information in the form of hypo- or hyper-methylation. Fortunately, bisulfite sequencing data suitable to investigate this possibility is readily available.

**Method:** We developed MethylSeqLogo, an extension of sequence logos which adds DNA methylation information to sequence logos. MethylSeqLogo includes new elements to indicate DNA methylation and under-represented dimers in each position of a set of aligned binding sites. Our method displays information from both DNA strands, and takes into account the sequence context (`CpG` or other) and genome region (promoter versus whole genome) appropriate to properly assess the expected background dimer frequency and level of methylation.

When designing MethylSeqLogo, we took care to preserve the usual sequence logo meaning of heights; in which the relative height of nucleotides within a column represents their proportion in the binding sites, while the absolute height of each column represents information (relative entropy) and the height of all columns added together represents total information.

**Results:** We present several figures illustrating the utility of using MethylSeqLogo to summarize data from `CpG` binding transcription factors. The logos show that unmethylated `CpG` binding sites are a feature of transcription factors such as MYC and ZBTB33, while some other `CpG` binding transcription factors, such as CEBPB, appear methylation neutral. We also compare MethylSeqLogo with two previously reported ways to create methylation aware sequence logos.

**Conclusions:** Our freely available software enables users to explore large-scale bisulfite and ChIP sequencing data sets — and in the process obtain publication quality figures.

## Background

Transcription Factors (TFs) are proteins which bind genomic DNA at specific sites (Transcription Factor Binding Sites: TFBSs) to regulate gene expression and thereby enable Eukaryotic cells to appropriately express genes according to: cell type, the cell cycle, the developmental stage of the organism, external conditions, etc. [1, 2, 3]. Moreover, perturbation of TF function plays major roles in the etiology of diseases such as cancer [4] and diabetes [5]. In humans these effects are realized by an ensemble of approximately 1600 TFs, each with distinct and often cell-type specific TFBSs [1, 6].

Given this importance and complexity, the study of TF function is both a long-standing and an on-going topic in molecular biology. One of the early successes in this endeavor was the invention of “sequence logos” [7], an effective way to visualize the position specific base preferences which partially characterize TFBSs. Sequence logos consist of a columns of the letters ({`A`, `C`, `G`, `T`} for a DNA motif) at each position, with the total column height of each position proportional to the information content of the distribution of bases in that position [7]. Their popularity attests to their utility in visually summarizing binding sites, which in turn facilitates communication (as figures in papers, etc.), and comparison between the binding preferences of distinct TFs. Indeed sequence logos have been extended in several ways; for example to improve the resolution of enriched/depleted components, e.g. Seq2logo [7] and EDlogo [8] or to show higher order sequence motifs [9] or inter-positional correlations in binding sites [10].

Sequence logos help biologists understand the sequence preference of TFs; but the local DNA sequence is only one factor determining binding site selection, and cannot explain cell type specific TFBS selection. Evidently, a more complete understanding of TF function requires the integration of local DNA sequence with epigenetic marks [11].

DNA methylation is particularly interesting because it can affect the binding of many transcription factors [12, 13, 14, 15, 16]; and is easily cast as DNA sequence information, since 5-methylcytosine can be viewed as a fifth DNA base [17, 18]. Moreover, technologies such as bisulfite sequencing can measure tissue specific genomewide DNA methylation levels at single-base resolution, and such data is already available for many cell types and conditions [19, 20].

Here we present MethylSeqLogo; a method which naturally extends classical sequence logos to visualize the methylation of a collection of TFBSs relative to an appropriate background. For user convenience we provide a software implementation prepackaged with methylation data for several cell lines from human, mouse, *Arabidopsis* and maize. The software also includes MethylScape, a companion method to MethylSeqLogo, which displays the methylation level of TFBS flanking regions.

## Visualization Method

Here we describe the design rationale and details of the MethylSeqLogo display; schematically presented in Figure 1.

**Figure 1.**
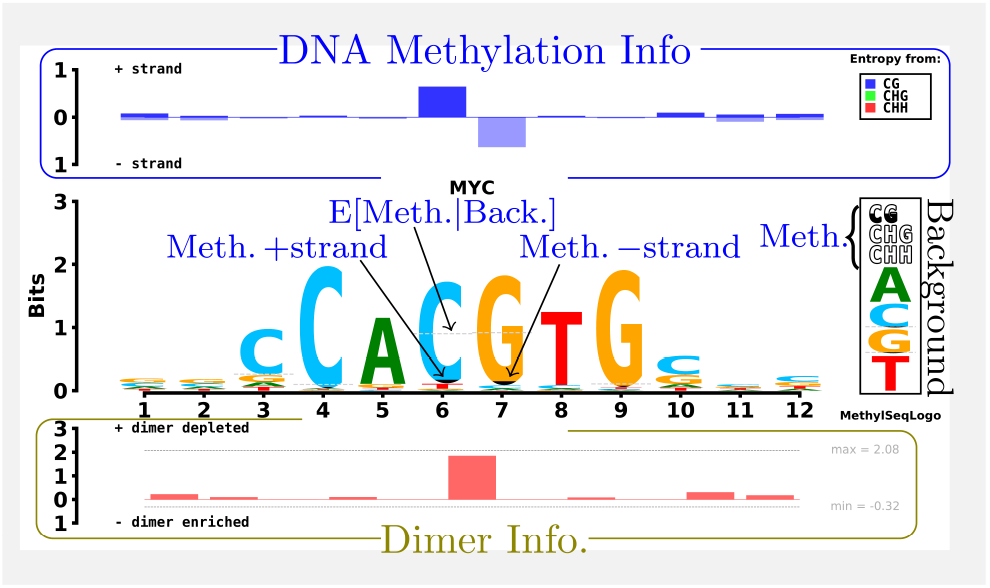
Design of MethylSeqLogo. Proportional shading of `C`’s and `G`’s indicates the methylation level of TFBS cytosines on the forward and reverse strands respectively; while a dashed line indicates the expected level of methylation based on the background distribution. The methylation key at lower right of the logo shows background methylation probabilities of `CG`, `CHG` and `CHH`, respectively; and the four single nucleotide background probabilities. The top track shows the relative entropy contributed by methylation in each context/strand combination, with information associated with cytosines on the reverse strand displayed downward. In the bottom track positive height indicating the presence of under-represented dimers (typically `CpG`), and negative height (not seen in this example) indicating the presence of over-represented dimers. For reference, the theoretical maximum and minimum possible dimer relative entropy contribution achievable for the given background are also shown.

### Design Goals

1. Keep the advantages of sequence logos; including familiarity.
2. For methylation, clearly display:
  - Strand (+*/−*) of the binding site
  - Trinucleotide context (`CG`, `CHG` or `CHH`)
  - Comparison relative to a background model
  - Dimer enrichment/depletion in the motif

We achieve the first goal by respecting two expectations viewers familiar with sequence logos will have: first, the relative height of an element (e.g. “`A`”) within a column represents the frequency of the corresponding element; and second, that the height of a column represents an information theoretic measure (relative entropy) of the degree to which that position in the binding sites differs from background [7]. We achieve the second goal by adding several intuitive elements to the plot:

- Partial shading of `C`’s and `G`’s
- Dashed line indicating expected methylation level
- Box at right showing background frequencies
- Context Colored Methylation info track at top
- Dimer enrichment/depletion info track at bottom

The height of the shading of `C`’s and `G`’s is proportional to the methylation level of cytosines on the forward and reverse strands respectively. In order to give users a clear image of hyper- or hypo-methylation, we added a dashed line showing the methylation level which would be expected based on the background distribution (taking the trinucleotide context {`CG, CHG, CHH`} in each binding site into account).

We designed a methylation info track showing (for each position in the binding site) the contribution of each context to the methylation information; and a box at right to show the background distribution of bases and methylation used for the relative entropy computation (Figure 1).

### Column Heights

This section describes how column height is determined for MethylSeqLogo’s three tracks in such a way that the total information in a set of binding sites can be estimated by visually adding up the height of all elements in a MethylSeqLogo display.

#### Column Heights Indicate Relative Entropy

Sequence logos often employ a background model fit to a set of background sequences, such as the whole genome or promoter regions etc. The background model is used to compute how “typical” the binding sequences are, with the idea that atypical binding site sequences should be emphasized visually (given taller column height) to reflect their statistical distance from background. For example, binding sites abundant in `C` and `G` should be emphasized more against an `AT`-rich background than against an `AT`-poor background. Quantitatively, the column heights are made proportional to the *relative entropy* ; also known as the *Kullback-Leibler directed divergence* [21], and equivalent to *information content* [22] under a uniform distribution background.

##### Sequence Background Models

In explaining the MethylSeqLogo sequence logo and dimer information tracks, we will refer to zero order Markov model and first order Markov model background models. Zero order models generate each nucleotide of DNA sequence independently, but first order models condition the nucleotide probabilities on the previous nucleotide.

##### Relative Entropy Formula

To facilitate describing the column heights of the MethylSeqLogo tracks in the following sections, we state the definition of relative entropy:

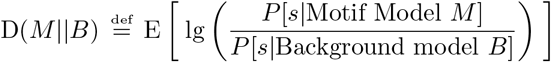

using lg to denote log_2_.

And the difference in relative entropy when employing background models **B**_1_ versus **B**_0_ is:

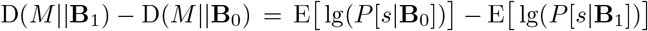

Where the expectation is the average over the individual binding site sequences *s* in a set of binding sites.

#### Sequence Logo Track Column Height

Standard sequence logos typically display columns with a height proportional to relative entropy using a PWM (Position Weight Matrix) based motif model which assigns distinct probabilities to the nucleotides {`A`,`C`,`G`,`T`} at each position but assumes independence between positions. A zero order Markov model, which also assumes positional independence, is usually employed as a background model. In this case the relative entropy of the binding sites is easily decomposed into a sum with one term for each position; and therefore can be conveniently displayed via the height of the column representing each position. MethylSeqLogo adopts these conventions for its sequence logo track.

#### Dimer Information Track

Although convenient, a zero order background model is unable to represent the striking (sometimes *>* 4x) depletion of `CpG` (relative to `CpC`, `GpC`, and `GpG` dinucleotides) in mammalian genomes. Admittedly, `CpG`’s are much less depleted in promoter regions, but there is still discrepancy between actual dimer frequencies versus what would be predicted by a zero order model. Therefore a first order model should provide a substantially more useful measure of how statistically distinct a set of binding sites is from background.

Given the potential size of this effect and the fact that methylation also happens at `CpG` dimers, we decided MethylSeqLogo should display information based on a first order Markov model background. We did not want to change the sequence logo track, so instead of directly displaying relative entropy against a first order Markov model, we chose to display the *difference* between that relative entropy and the zero order background relative entropy in a separate track. Fortunately, this difference can easily be decomposed into the sum of a set of terms; one term for each pair of adjacent positions (see supplementary text for a mathematical derivation). Since these terms represent pairs of adjacent nucleotide positions, MethylSeqLogo displays them as vertical bars between the two positions. In theory, column heights in this track can be negative if the binding sites contain many over-represented dimers (for example homodimers `XpX` may be somewhat over-represented).

#### Methylation Track Column Height

Hyper- or hypo-methylation of TF bindings sites (relative to a background) may help distinguish those binding sites from that background. To allow users to see this effect, MethylSeqLogo presents a methylation information track above the main sequence logo track. Informally, the height of bars in the methylation information track represent the amount of additional surprise experienced when observing the methylation value at position *i* from one of the TFBSs; *after* having observed the primary sequences, since that information is already accounted for in the other tracks. The propensity of genomic cytosines to be methylated differs strongly depending on the following base or two (i.e. `CG`, `CHG`, or `CHH` trinucleotide context), so we separate these cases in our computation. For a background distribution these three cases are enough; while for binding sites, position and strand must also be considered. Thus altogether we separate the methylation data for each position in a collection of TFBSs into 6 strand specific contexts: 3 trinucleotide contexts *×* 2 strands (Supplementary Figure 2).

Formally, let *P*_context | *i*_ denote the probability that a binding site will have a cytosine matching the given context at position *i* and *P*_*m* | context,*i*_ denote the probability that such a cytosine will be methylated or not; while *P*_*m* | context,BG_ denotes the background probability of a cytosine in that given context being methylated or not. We can write the contribution of methylation information to the height of column *i* as:

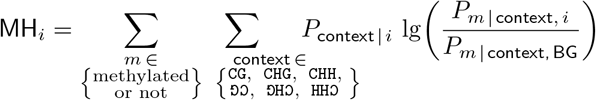

Note that relative entropy is inherently robust to small sample estimation error in *P*_*m* | context, *i*_ since it includes a multiplicative term *P*_context | *i*_ in the contribution of that context to column height. Thus rare contexts cannot make large contributions to column height.

## Data and Software

### DNA Methylation and TFBS data

MethylSeqLogo requires binding sites and methylation information, preferably specific to a given tissue or cell-type. To gather this information we built a computational pipeline to process ChIP-seq data for TFBSs and WGBS to calculate the methylation probability of each position in the aligned TFBSs, as well as the background probabilities (Supplementary Figure 1).

#### Whole Genome Bisulfite Sequencing Data

We downloaded Human reference genome GRCh37 (hg19) and GRCh38 (hg38) from the Illumina iGenomes website and Human WGBS (Whole Genome Bisulfite Sequencing) from the ENCODE [23] website. The figures in this publication reflect data from ENCODE IDs: (`086MMC`, `379ZXG`, `417VRB`, `524BMX`, `601NBW`, `918PML`) and (`030LDK`, `086KJC`, `300GSM`, `390OZB`, `624VFJ`, `847OWL`) for H1-hESC and HepG2 cell lines respectively (all IDs start with `ENCFF`).

We merged the methylation calling BED files of two replicates for each cell type, by averaging the methylation levels of cytosine sites (on either strand) with read depth greater than four.

#### Methylation of TFBSs

We collected TFBS coordinates from the JASPAR database [24]; and tissue-specific ChIP-seq data from ReMap [25] for TFs (EGR1, MYC, SP1, USF1 and ZBTB33) to generate the figures in this text, and CEBPB for a supplementary text figure. To obtain tissue-specific TFBS coordinates, we used the `bedtools intersect` function. Based on those coordinates, one can generate the intermediate input files needed by MethylSeqLogo to generate MethylSeqLogo images.

### Promoter Regions

Promoter regions have special significance for most transcription factors, but the distribution of both `CpG`’s and their methylation differs sharply between promoters regions and the genome as a whole. Thus we provide predefined promoter regions defined as 1000bp upstream to 200bp downstream of annotated major transcription start sites [26].

### MethylSeqLogo Program

We provide an open source implementation of the MethylSeqLogo visualization method and a companion program MethylScape described in below.

MethylSeqLogo comes with precomputed probability models for the examples discussed in this paper and many other tissues that have published WGBS data. Users can also calculate the methylation probabilities from their own WGBS datasets with a script provided in the MethylSeqLogo package and generate logos of that data.

## Example MethylSeqLogos

### MYC Binding Sites

MYC transcription factors (data show here is for c-Myc) are oncogenic transcription factors that bind DNA as a heterodimer with MAX [27]. Figure 2 (left) shows shows MethylSeqLogo’s of MYC using data from H1-hESC cells (numerical data shown in Supplementary Table 1). From these images it is apparent that when looking at the entire genome, MYC binding sites are statistically characterized by hypomethylation and the occurrence of the under-represented dimer `CpG`. On the other hand, the promoter region based MethylSeqLogo’s (Figure 2 (top)) shows greatly reduced information from hypo-methylation and `CpG`; but it is “cleaner” in the sense that the methylation information is concentrated at positions 6 and 7, consistent with reports that methylation in the center `CpG` site of MYC binding sites reduces binding efficacy [28].

**Figure 2.**
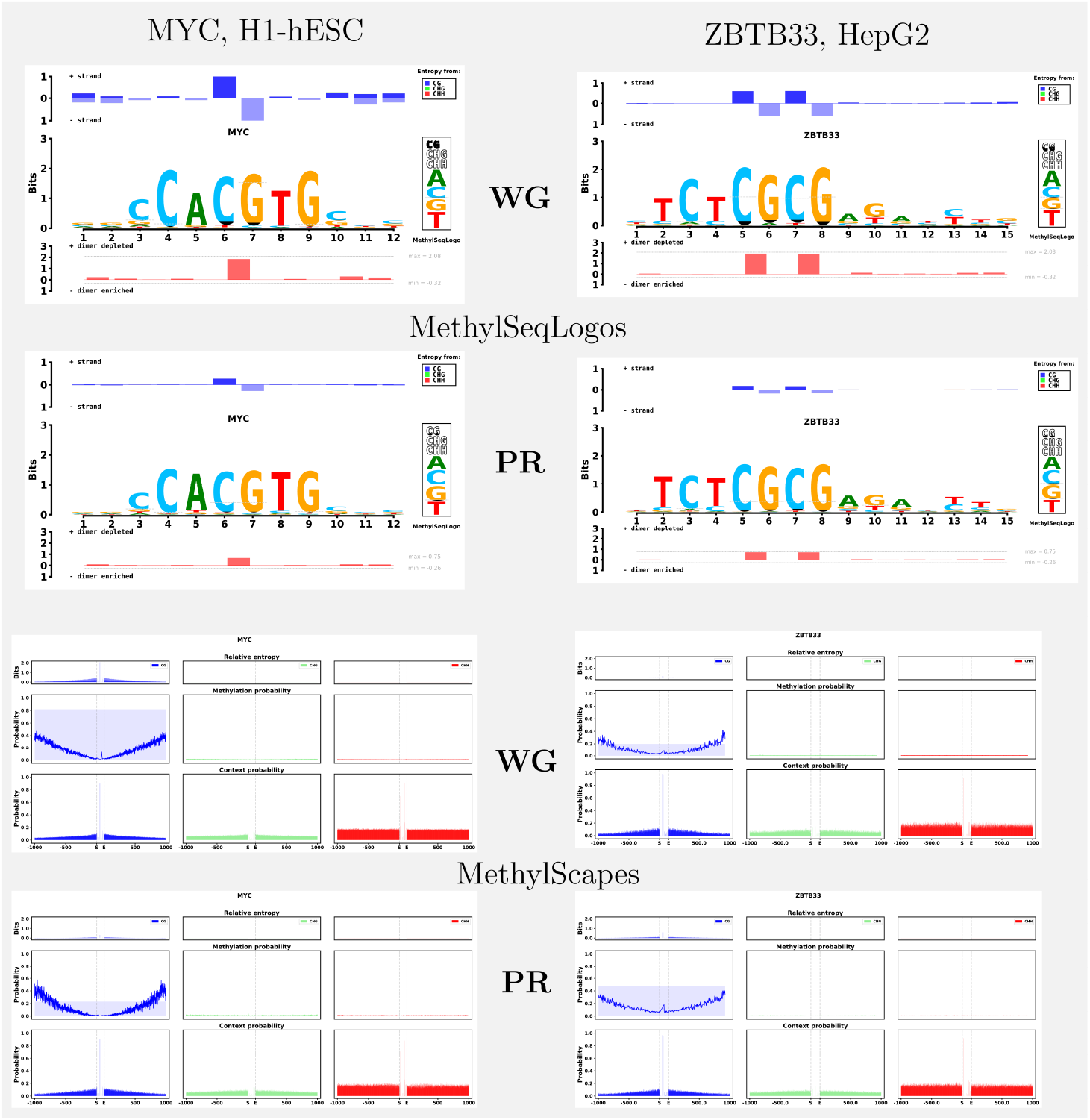
MethylSeqLogos (top) and MethylScape(s) (bottom) of c-Myc binding sites in H1-hESC cells (left) and ZBTB33 binding sites in HepG2 cells (right). Logos in rows marked with **WG** show information for all binding sites relative to a whole genome (WG) background model, while logos in rows marked with **PR** show information for promoter region binding sites relative to a promoter region (PR) background model. The three columns in a MethylScape logo represent the contexts: `CpG`, `CHG`, and `CHH`; with faint background color in the middle row representing the background model methylation probability for each respective context.

### ZBTB33 Binding sites

Figure 2 (right) shows MethylSeqLogos for ZBTB33 in HepG2 cells. ZBTB33, also named Kaiso [29], is a homodimeric transcription factor associated with several types of cancer [30]. ZBTB33 has been reported to bind methylated `CpG`’s and the sequence motif TCCTGCNA [31], especially TCTCGCGAGA [32]; with *in vitro* data indicating a much higher affinity for this motif when methylated. Comparing ChIP-Seq and bisulfite sequencing data, Blattler et al. [33] were able to confirm the TCTCGCGAGA motif, but found that very few ZBTB33 binding sites are methylated *in vivo*. The visual impression given by MethylSeqLogo is in line with their conclusions.

### Contrasting TF binding motifs

Transcription factors can be grouped by structural features of their DNA-binding domains. Often TFs with the same type of DNA-binding domains will bind to similar DNA sequences, which are sometimes called *response elements*. For example, an E-box (enhancer element) is a response element with palindromic general pattern `CANNTG` (`N` denotes any base) and canonical sequence `CACGTG`.

MYC and USF1 both have bHLH (basic helix-loop-helix) DNA-binding domains which bind to canonical E-box response elements. Comparing the methylation track of their MethylSeqLogos in figure 3 (left), USF1 appears more tolerant of methylation of the cytosines in the central `CpG`. Interestingly, comparing the sequence logo tracks, one can see that USF1 binding sites also exhibit more frequent substitution of 5-methyluracil (i.e. thymine) for cytosine as well. This example illustrates the utility of MethylSeqLogo in simultaneously comparing the primary sequence and methylation preferences of DNA binding motifs. Figure 3 (right) shows another example, comparing the GC-box element transcription factors SP1 and EGR1.

**Figure 3.**
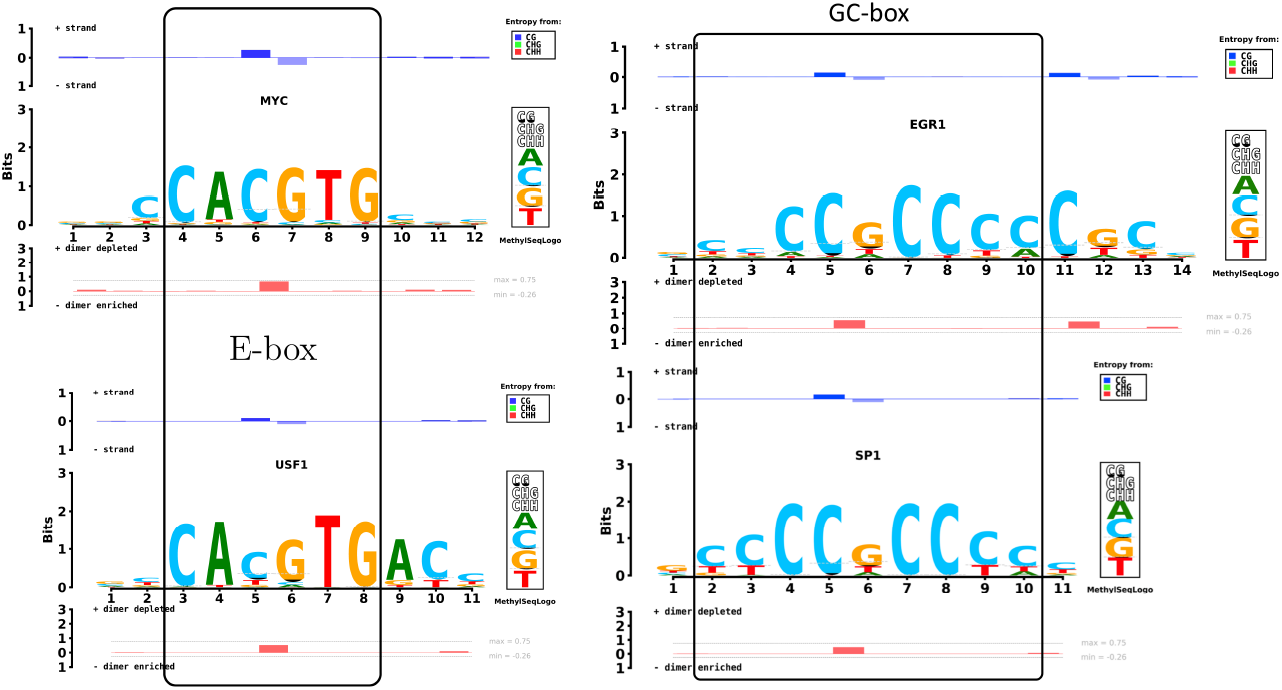
MethylSeqLogo facilitates comparison of the DNA methylation of transcription factors with similar binding preferences. The E-box elements binding TFs MYC and USF1 (at left) and the the GC-box elements binding TFs SP1 and EGR1 (at right) are compared using promoter region binding sites and background model. Data from H1-hESC cells.

### MethylScape shows methylation relative entropy in a wider window

The cytosine methylation levels around TFBSs may relate to TF binding [34]. Therefore we developed MethylScape, a companion program to MethylSeqLogo, that can display methylation entropy, methylation probability and context probability trends around TFBSs. Figure 2 (bottom) shows MethylScape plots of MYC and ZBTB33 whole genome and promoter region binding sites. Compared to the flanking regions, MYC binding sites are hypo-methylated, in both promoter and whole genome (middle MethylScape panel), even though some `CpG`’s can be seen near the binding sites (bottom MethylScape panel, left columns). Since hypo-methylation is somewhat less surprising in promoter regions, the `CG` in the center of the MYC binding site is more prominent in the whole genome MethylScape than in the promoter regions MethylScape.

### Related Visualization Tools

Some other methods have been previously been proposed to extend sequence logos to include DNA methylation information. MeDReaders [35] is a database summarizing methylation level with TFBS coordinates. MethMotif [36] is a database organizing tissue-specific data. Both of these resources provide methylation aware sequence logos for the convenience of their users. While Meth-eLogo [37] extends affinity (energy) sequence logos to include DNA methylation. The visual design of these tools is completely different than MethylSeqLogo (see supplementary material for a comparison).

## Discussion

### Caveats

When viewing MethylSeqLogo’s one must keep in mind the choice of background. In particular, since TFs tend to bind promoter regions, and promoter regions tend to be hypo-methylated; under a whole genome background, MethylSeqLogo’s will tend to show some amount of hypo-methylation for any `CpG` containing binding site. This effect can be seen in the logos shown in Figure 2 — under a whole genome background the MYC logos shows methylation information distributed across many positions, but under a promoter region background only some of the methylation information in the central binding motif `CpG` remains. In a narrow sense both logos faithfully depict statistical differences between binding sites and the respective background; but in terms of the impression given, a whole genome background may seem to exaggerate the importance of methylation.

On the other hand, the MethylSeqLogo’s displayed here may also understate the importance of methylation on TF binding. The methylation and TF binding data used here are the average of many cells from two samples (of the same cell line, but not the same cells), so if TF binding and methylation vary between cells or samples, the correlation between them will be under-estimated. Measurement noise (unless systematically biased) will also tend to decrease correlations. Therefore the correlations presented in the logos here may be reduced in magnitude.

### Future Work

#### Tailored background models

The particular definition of promoter regions we used here seems to work well but may not always be the most appropriate. Certainly more choices could be offered, perhaps: core promoter, extended promoter, promoter + known enhancer regions, etc. Going one step further, background regions could be tailored for a given set of TFBSs, by using regions within some distance (say 50bp) of each binding site. Ideally this would be done independently for each binding site (so that a genome position near *x* binding sites would be include *x* times in the background model statistics). Thus ensuring the statistical differences depicted in MethylSeqLogo logos would be due to the binding sites (or at most their immediately flanking bases), rather than larger scale trends in methylation and/or `CpG` frequency across the genome.

#### Displaying More Information

*Other Cytosine Modifications:* 5-hydroxymethyl cytosine (5hmC) is an intermediate in the demethylation pathway from `5mC` to unmethylated cytosine [38]. These three forms of cytosine have distinct chemical structures and may provide distinct binding affinities for DNA binding proteins [39]. But the data presented in this manuscript lumps `5mC` and `5hmC` together, as standard bisulfite sequencing cannot distinguish between them [40, 41]. Fortunately, data specific for `5hmC` is becoming available [42] and extending MethylSeqLogo to visualize that data should be relatively manageable; perhaps modeling the distinct between `5mC` and `5hmC` as an additional piece of information gained after learning that a cytosine is modified in some way (is either `5mC` and `5hmC`). Conveniently, like `5mC`, `5hmC` occurs primarily in `CpG` context [42]; which MethylSeqLogo already treats specially.

*Other Epigenetic Information:* We briefly considered the display of other forms of DNA modification — or, more ambitiously, histone modification. We are aware of one attempt to display histone modification in a sequence logo type display, but only at a very broad resolution of introns, exons, etc. [8]. Indeed, since histone marks are not associated with single DNA residues, and in general may be positioned differently at each binding site of a TF, it is not clear where a ‘column’ in a histone mark sequence logo should begin and end. Thus sequence logos may not turn out to be a better way of visualize histone modification than approaches such as juxtaposing [43] or averaging [44] heatmaps or ‘wiggles’. One concept from sequence logos which might be applicable would be to try making the area of histone mark wiggles proportional to some measure of their information relative to a background model. In any case, visualizing histone modification is beyond the scope of this work.

#### Dimers track information could be displayed as letters

Currently the MethylSeqLogo displays the dimer track simply as bars of indicating total column height. One could imagine using sequence-logo-like letters in this track instead of bars. So for example, “`CG`” could be drawn with height proportional to the contribution of `CpG` to the dimer information. In the examples shown in this manuscript, `CpG` is in fact responsible for the bulk of the information in the dimer track, so if rendered as “`CG`” it should be tall enough to be legible in some cases. Nevertheless, when designing the display we felt that a lettered dimer track would overall be more distracting than informative. The idea might be worth exploring in the future however, especially since the idea of a background model based dimer track is not specific to methylation and could be added to any sequence logo, even protein sequence logos.

#### Higher Order Background Models

Finally we note that in principle a “trimer information track” (or even higher order tracks) could be added to the display, with each level showing the change in relative entropy resulting in incrementing the background model order. This approach might make sense in applications where the background sequence has significantly under/over-represented trimers (e.g. DNA sequences coding for proteins).

### Conclusions

Sequence logos are the method of choice to visualize the nature and strength of the local primary DNA sequence contribution to TFBS selection. DNA methylation also contributes significantly to binding site selection for some transcription factors and DNA methylation data is conveniently analogous to the primary sequence data used for traditional sequence logos. Thus it is natural and desirable to extend sequence logos to include DNA methylation. We believe MethylSeqLogo has accomplished this and will prove useful.

MethylSeqLogo comes with precomputed probability models for many tissues that have published WGBS data. Users can also calculate the methylation probabilities of their own WGBS dataset with a script provided in the MethylSeqLogo package and plot on the basis of that background. Complementing MethylSeqLogo, MethylScape gives a wider view around TFBSs.

## Acknowledgements

We thank Dr. Martin Frith, Dr. Toshikazu Ushijima, Dr. Naoko Ida, Dr. Tony Kuo and Dr. Matteo Pellegrini for kindly providing comments on the manuscript.

## Abbreviations

H1-hESC: Human Embryonic Stem Cell line H1
TF: Transcription Factor
TFBS: Transcription Factor Binding Sites
WGBS: Whole Genome Bisulfite Sequencing

## Availability of data and materials

Source code: https://gitlab.com/fmhsu0114/methylseqlogov2

Gallery: https://feimanh.shinyapps.io/MethylSeqLogo

Data: The figures in this manuscript were generated based on publicly available data. The sources are listed in the main text.

## Ethics approval and consent to participate

Not applicable.

## Consent for publication

Not applicable.

## Authors’ contributions

P.H. conceived of the study. FM.H. and P.H. designed the display. FM.H. implemented the software and selected the test cases shown. FM.H. and P.H. wrote the manuscript.

## Additional Files

Additional file 1 — Supplemental Text

A pdf document provides additional figures and test giving details of our method and a comparison to related methods.

