## Supplemental Text for "MethylSeqLogo: DNA methylation smart sequence logos"

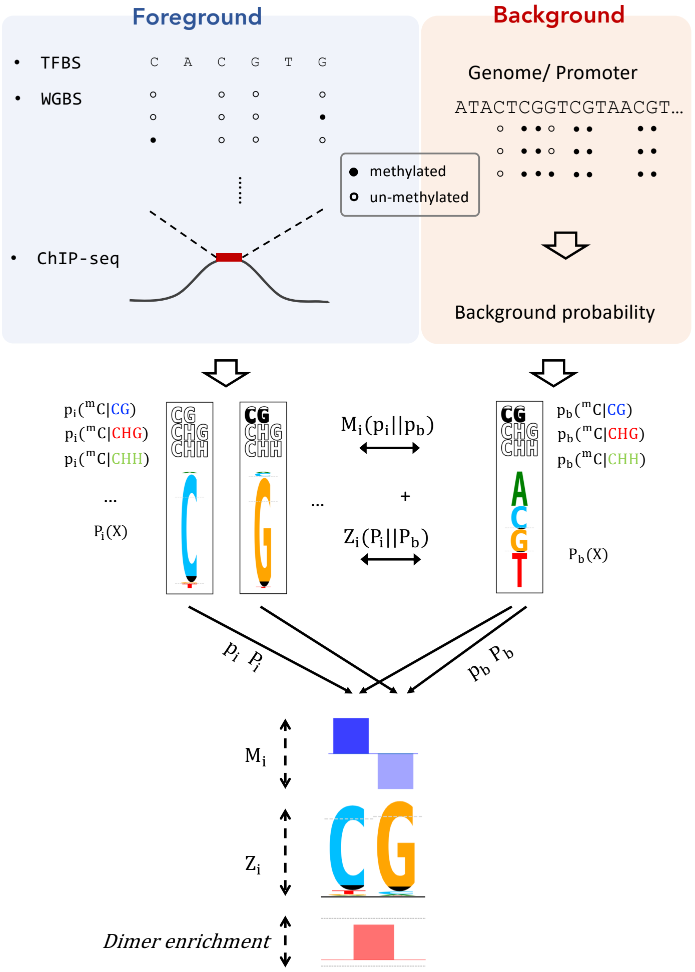


Figure 1: MethylSeqLogo pipeline. Tissue-specific TFBSs are acquired by incorporating ChIP-seq data. Cytosine methylation probabilities inside TFBSs and the genome are from WGBS. $P(X)$ represents the foreground (motif) and background (genome) probability of A, C, G, T, while p($^m$C|CG), p($^m$C|CHG) and p($^m$C|CHH) represent the conditional probability of $^m$C in each context. The sum of $Z_i$ and $M_i$ is used as the column height of position $i$ inside a TF binding motif. The black region of letters C and G represent methylated cytosine on the forward and reverse strand respectively.

### Methods (supplemental)

#### Workflow

Figure 1 (shown on the previous page) outlines the workflow used to produce the logos in this publication.

#### Cytosine Contexts

Figure 2 illustrates the contexts of genomic cystosines treated distinctly in this work.

| | |
|---|---|
| Forward Strand | `taCCGACGTGACCtg` |
| Methylation Values | `--377-77-2-23--` |
| Opposite Strand | `atGGCTGCACTGGac` |

Figure 2: Various contexts of cytosines are illustrated. The top and bottom sequences represent the forward and opposite strand of an aligned TFBS. Residues on the opposite strand are drawn upside-down. The middle track represents a simplified, single digit depiction of the methylation values ("2" represents 20%, "7" represents 70%, etc.) associated with the cytosines in the TFBS, with values associated with a cytosine on the opposite strand drawn upside-down. Cytosines and their associated methylation values are color coded by {`CG`,`CHG`,`CHH`} trinucleotide context. Upper-case letters indicate the width of the binding motif, with two flanking bases on each side to determine cytosine trinucleotide context.

#### Methylation Probability Estimation

In this section we describe how we estimate the probability that cytosines are methylated given their context. First we use bisulfite sequencing reads aligned to the reference genome to obtain an estimate of the proportion of the genomic DNA molecules for which methylation protected the cytosine at position $j$ from conversion to uracil (producing a `T` in the sequence data). We use $v_j$ to denote this estimate and refer to it as the *methylation value* (these methylation values can themselves be viewed as probabilities, but we use the term "value" here to help distinguish them from other probabilities defined below).

To precisely define the methylation values, let $\mathbf{R}_j$ denote the set of (uniquely mapped) reads aligning a base (e.g. not a gap) to position $j$; reverse complementing those reads as necessary so that they all align to the reference strand. Using $\mathbf{R}_j^{\mathrm{C/C}}$ to denote the case of `C` aligned to `C` (i.e. no evidence of bisulfite conversion of the genomic cytosine) we define $v_j$ as:

$$v_j = \frac{|\mathbf{R}_j^{\mathrm{C/C}}|}{|\mathbf{R}_j|} \quad \text{if } |\mathbf{R}_j| \geq \theta$$

Where $\theta$ is minimum read coverage threshold, which we sent to 4, following usual practice for bisulfite sequencing data Stroud *et al.* (2013). We discard sites with coverage less than $\theta$ in subsequent computations.

Next we estimate the background methylation probabilities separately for each of the three contexts {CG, CHG, CHH}, by simply averaging the defined methylation values (the $v_j$'s) of background sequence cytosines in each context respectively. Even when requiring a minimum coverage of 4, these probability estimates are typically based on many ($\gg 1000$) genomic cytosines and therefore do not require carefully set priors.

In contrast, the sequences observed at certain positions of TFBSs may be highly constrained, leading to small samples for some combinations of TFBS strand, position and trinucleotide context. Fortunately relative entropy is inherently robust to inaccurate estimates of rare events, since $\lim_{p \to 0} p \lg p = 0$. So accurate estimation of the methylation level of combinations which only appear a few times is not critical. Nevertheless we employ priors based on the (trinucleotide context specific) background methylation probabilities to obtain reasonable estimates even for rare combinations.

$$P_{\mathrm{meth} \,|\, \mathsf{context},\, i} \stackrel{\mathrm{def}}{=} \frac{P_{\mathrm{meth} \,|\, \mathsf{context,BG}} + n_{\mathsf{context},\, i} \cdot \overline{v} \,\big|\, \mathsf{context},\, i}{1 + n_{\mathsf{context},\, i}}$$

$$\mathsf{context} \in \left\{ \begin{matrix} \mathtt{CG, CHG, CHH,} \\ \mathtt{⅁Ɔ, ⅁HƆ, HHƆ} \end{matrix} \right\},\ 0 \le i < w$$

| Notation | Description |
|---|---|
| $P_{\mathrm{meth} \,\vert\, \mathsf{context,BG}}$ | The background probability of methylation by context. |
| $n_{\mathsf{context},\, i}$ | The number of cytosines (with sufficient read coverage) of the given context occurring at position $i$ in the set of TFBS. |
| $\overline{v} \,\vert\, \mathsf{context},i$ | The average methylation values of those cytosines. |

Figure 3: Position specific, context sensitive, methylation probability defined.

The equation shown in figure 3 for $P_{\mathrm{meth} \,|\, \mathsf{context},\, i}$ reduces to $P_{\mathrm{meth} \,|\, \mathsf{context,BG}}$ when there is no data for the given context; and to a Jeffreys prior when $P_{\mathrm{meth} \,|\, \mathsf{context,BG}}$ happens to equal $1/2$. Naturally, the probability of cytosines in any context being unmethylated is simply set to one minus the estimated probability of methylation.

Thus altogether we separate the methylation data for each position in a collection of TFBSs into 6 strand specific contexts: 3 trinucleotide contexts $\times$ 2 strands, which we denote as {CG, CHG, CHH} and {⅁Ɔ, ⅁HƆ, HHƆ} for the forward and reverse strand respectively (Figure 2). Note that these strand specific contexts are mutually exclusive and together cover every possible way a cytosine can occur.

#### Derivation of Dimer Track Column Height Computation

The dimer information track column heights are designed to be proportional to the difference in relative entropy obtained when substituting a first order Markov model for a zero order Markov model (both models trained on the same background sequences). Here, we derive the decomposition of this difference into a sum of terms, one term for each pair of adjacent nucleotide positions.

Table 1: MYC methylation probabilities, relative entropy and column heights corresponding to the MethylSeqLogo shown in Figure 2 of the main manuscript. Columns 2–7 show the contribution to total relative entropy due to methylation in each of the six strand specific contexts. $M_i$ denotes the methylation relative entropy, $Z_i$ the canonical {A,C,G,T} based relative entropy, and $M_i + Z_i$ the MethylSeqLogo column height. Numbers of interest are highlighted in blue.

| **Promoter Regions Background:** (CG, CHG, CHH)= (0.23, 0.01, 0.01) | | | | | | | | | |
|---|---|---|---|---|---|---|---|---|---|
| $i$ | CG | ᘓƆ | CHG | ᘓHƆ | CHH | HHƆ | $M_i$ | $Z_i$ | $M_i + Z_i$ |
| 1 | 0.04 | 0.02 | 0.00 | 0.00 | 0.00 | 0.00 | 0.07 | 0.09 | 0.16 |
| 2 | 0.01 | 0.04 | 0.00 | 0.00 | 0.00 | 0.00 | 0.05 | 0.10 | 0.15 |
| 3 | 0.00 | 0.01 | 0.00 | 0.00 | 0.00 | 0.00 | 0.02 | 0.78 | 0.79 |
| 4 | 0.01 | 0.00 | 0.00 | 0.00 | 0.00 | 0.00 | 0.01 | 1.65 | 1.67 |
| 5 | 0.00 | 0.01 | 0.00 | 0.00 | 0.00 | 0.00 | 0.01 | 1.48 | 1.49 |
| 6 | 0.27 | 0.00 | 0.00 | 0.00 | 0.00 | 0.00 | 0.27 | 1.56 | 1.83 |
| 7 | 0.00 | 0.28 | 0.00 | 0.00 | 0.00 | 0.00 | 0.28 | 1.60 | 1.88 |
| 8 | 0.01 | 0.00 | 0.00 | 0.00 | 0.00 | 0.00 | 0.01 | 1.53 | 1.54 |
| 9 | 0.00 | 0.01 | 0.00 | 0.00 | 0.00 | 0.00 | 0.01 | 1.58 | 1.60 |
| 10 | 0.04 | 0.00 | 0.00 | 0.00 | 0.00 | 0.00 | 0.04 | 0.28 | 0.32 |
| 11 | 0.03 | 0.04 | 0.00 | 0.00 | 0.00 | 0.00 | 0.07 | 0.09 | 0.16 |
| 12 | 0.03 | 0.03 | 0.00 | 0.00 | 0.00 | 0.00 | 0.07 | 0.15 | 0.22 |
| **Whole Genome Background:** (CG, CHG, CHH)= (0.82, 0.02, 0.01) | | | | | | | | | |
| 1 | 0.22 | 0.18 | 0.00 | 0.00 | 0.00 | 0.00 | 0.40 | 0.19 | 0.59 |
| 2 | 0.08 | 0.21 | 0.00 | 0.00 | 0.00 | 0.00 | 0.30 | 0.19 | 0.49 |
| 3 | 0.01 | 0.08 | 0.00 | 0.00 | 0.00 | 0.00 | 0.10 | 1.03 | 1.13 |
| 4 | 0.08 | 0.01 | 0.00 | 0.00 | 0.00 | 0.00 | 0.10 | 2.10 | 2.20 |
| 5 | 0.00 | 0.08 | 0.00 | 0.00 | 0.00 | 0.00 | 0.08 | 1.27 | 1.35 |
| 6 | 1.80 | 0.00 | 0.00 | 0.00 | 0.00 | 0.00 | 1.80 | 1.87 | 3.67 |
| 7 | 0.00 | 1.84 | 0.00 | 0.00 | 0.00 | 0.00 | 1.84 | 1.96 | 3.80 |
| 8 | 0.07 | 0.00 | 0.00 | 0.00 | 0.00 | 0.00 | 0.07 | 1.36 | 1.43 |
| 9 | 0.00 | 0.07 | 0.00 | 0.00 | 0.00 | 0.00 | 0.07 | 2.07 | 2.15 |
| 10 | 0.26 | 0.00 | 0.00 | 0.00 | 0.00 | 0.00 | 0.26 | 0.60 | 0.86 |
| 11 | 0.19 | 0.28 | 0.00 | 0.00 | 0.00 | 0.00 | 0.47 | 0.11 | 0.58 |
| 12 | 0.21 | 0.18 | 0.00 | 0.00 | 0.00 | 0.00 | 0.40 | 0.26 | 0.67 |

The relative entropy we use is defined as:

$$\text{Relative Entropy} \stackrel{\text{def}}{=} \mathrm{E}\left[\ \lg\left(\frac{P[s|\text{Motif Model}]}{P[s|\text{background model}]}\right)\ \right]$$
$$= \mathrm{E}\Big[\ \lg(P[s|\text{Motif Model}]) - \lg(P[s|\text{background model}])\ \Big]$$
$$= \mathrm{E}\Big[\ \lg(P[s|\text{Motif Model}])\ \Big] - \mathrm{E}\Big[\ \lg(P[s|\text{background model}])\ \Big]$$

Where the expectation is the average over individual binding site sequences $s$ in a set of binding sites $S$.

From this definition we can see that (with a constant motif model) the difference in relative entropy when switching from a zero to first order Markov model for the background becomes:

$$\text{Rel. entropy diff} = \mathrm{E}\Big[\lg(P[s|\mathbf{B}_0]) - \lg(P[s|\mathbf{B}_1])\Big] = \mathrm{E}\left[\lg\left(\frac{P[s|\mathbf{B}_0]}{P[s|\mathbf{B}_1]}\right)\right]$$

Where $\mathbf{B}_0$ and $\mathbf{B}_1$ denote zero and first order Markov models reflecting background frequencies of nucleotides and dinucleotides respectively. Repeating their definitions here:

$$P[s|\mathbf{B}_0] = \prod_{i=1}^{w} P[s_i|\mathbf{B}_0]$$

$$P[s|\mathbf{B}_1] = P[s_1|\mathbf{B}_0] \prod_{j=2}^{w} P[s_j|s_{j-1}, \mathbf{B}_1]$$

Where $s = s_1 \ldots s_w$ denotes a binding site sequence, $P[s_j|s_{j-1}, \mathbf{B}_1]$ the probability of base $s_j$ given the previous base and the probabilities held in $\mathbf{B}_1$. To simplify the visual presentation of our derivations, we define $i \stackrel{\text{def}}{=} j-1$, so $s_{j-1}s_j$ can be written as $s_i s_j$. Yielding,

$$P[S|\mathbf{B}_1] = P[s_1|\mathbf{B}_0] \prod_{j=2}^{w} P[s_j|s_{j-1}, \mathbf{B}_1] \stackrel{\text{def}}{=} P[s_1|\mathbf{B}_0] \prod_{j=2}^{w} P[s_j|s_i, \mathbf{B}_1]$$

Typically sequence logo heights are computed relative to $\mathbf{B}_0$, but we would like to extend this by adding a dinucleotide track showing the extra relative entropy gained by considering the background distribution of dinucleotides. In mathematical terms we seek a quantity $r_j$ such that:

$$P[s|\mathbf{B}_1] = P[s|\mathbf{B}_0] \prod_{j=2}^{w} r_j \tag{1}$$

Where $\lg(r_j)$ represents the extra relative entropy resulting from considering the dinucleotide $s_i s_j$ together after considering $s_i$ and $s_j$ separated (reminder: we use $s_i$ to denote $s_{j-1}$).

We should be able to derive $r_j$ from equation 1, but instead we immediately guess at the answer and then show it is correct. Based on standard measures of correlation, our guess at $r_j$ is:

$$r_j := \frac{P[s_i s_j|\mathbf{B}_1]}{P[s_i s_j|\mathbf{B}_0]} = \frac{P[s_i s_j|\mathbf{B}_1]}{P[s_i|\mathbf{B}_0]\,P[s_j|\mathbf{B}_0]}$$

Substituting this equation into equation 1 we obtain:

$$\begin{aligned}
P[s|\mathbf{B}_1] = \quad & P[s|\mathbf{B}_0] \quad \prod_{j=2}^{w} \frac{P[s_i s_j|\mathbf{B}_1]}{P[s_i|\mathbf{B}_0]\,P[s_j|\mathbf{B}_0]} \\
& \prod_{j=1}^{w} P[s_j|\mathbf{B}_0] \quad \prod_{j=2}^{w} \frac{P[s_i s_j|\mathbf{B}_1]}{P[s_i|\mathbf{B}_0]\,P[s_j|\mathbf{B}_0]} \\
& P[s_1|\mathbf{B}_0] \prod_{j=2}^{w} P[s_j|\mathbf{B}_0] \prod_{j=2}^{w} \frac{P[s_i s_j|\mathbf{B}_1]}{P[s_i|\mathbf{B}_0]\,P[s_j|\mathbf{B}_0]} \\
& P[s_1|\mathbf{B}_0] \prod_{j=2}^{w} \frac{P[s_j|\mathbf{B}_0]\,P[s_i s_j|\mathbf{B}_1]}{P[s_i|\mathbf{B}_0]\,P[s_j|\mathbf{B}_0]} \\
& P[s_1|\mathbf{B}_0] \prod_{j=2}^{w} \frac{P[s_i s_j|\mathbf{B}_1]}{P[s_i|\mathbf{B}_0]} \\
& P[s_1|\mathbf{B}_0] \prod_{j=2}^{w} \frac{P[s_i s_j|\mathbf{B}_1]}{P[s_i|\mathbf{B}_1]} \\
& P[s_1|\mathbf{B}_0] \prod_{j=2}^{w} P[s_i s_j|s_i, \mathbf{B}_1] \;=\; P[s_0|\mathbf{B}_1] \prod_{j=2}^{w} P[s_j|s_i, \mathbf{B}_1] = P[S|\mathbf{B}_1] \checkmark
\end{aligned}$$

Where $P[s_i|\mathbf{B}_1] \equiv P[s_i|\mathbf{B}_0]$ because the first order Markov model background model reduces to a zero order model when generating only one nucleotide.

The last line follows from the definition of conditional probability $P[j|i] \stackrel{\text{def}}{=} \frac{P[i,j]}{P[i]}$, by comparing that to $P[s_i s_j|s_i]$ and noting this could equivalently be written as simply $P[s_j|s_i]$ since $s_i$ is given.

### Results (supplemental)

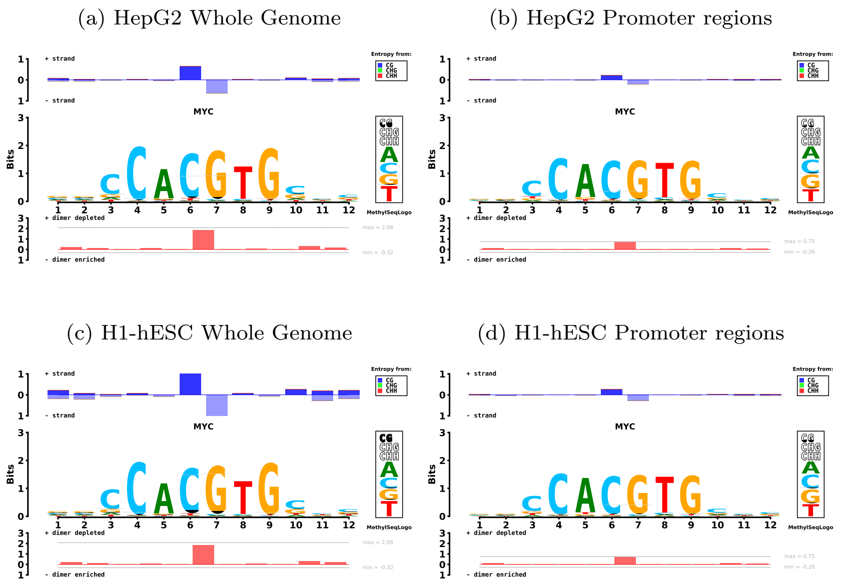


Figure 4: MYC MethylSeqLogo's for Whole Genome (at left) and Promoter Regions (at right) in HepG2 (top) and H1-hESC (bottom) tissue types.

#### Tissue-specific TFBS Logos

In contrast to the nearly constant primary DNA sequence, genome DNA methylation varies widely depending on cell type and developmental stage. In this section we explore the use of MethylSeqLogo to compare the DNA methylation of TFBSs in different cell types.

Figure 4 contrasts the MethylSeqLogo's for MYC binding in two cells lines: HepG2, a cell line with relatively low methylation; and H1-hESC. Note that both background methylation probabilities and the binding sites themselves differ between tissue types. As might be expected, the hypo-methylated central `CpG` in the MYC binding site is least prominent (i.e. least surprising) in the relatively unmethylated HepG2 promoter regions.

A contrary case is the CCAAT/Enhancer-binding protein CEBPB, a transcription factor for which various forms of modified cytosine (not only 5-methylcytosine, but also 5-hydroxymethylcytosine, 5-formylcytosine and 5-carboxylcytosine) are reported to have mixed effects on binding *in vitro* (Sayeed *et al.*, 2015). Unlike MYC, the MethylSeqLogo for CEBPB shows hardly any difference in methylation between binding sites and background for either whole genome or promoter regions (Figure 5 (a)), indicating that the methylation level of CEBPB binding sites tracks the background of the genomic regions they occur in.

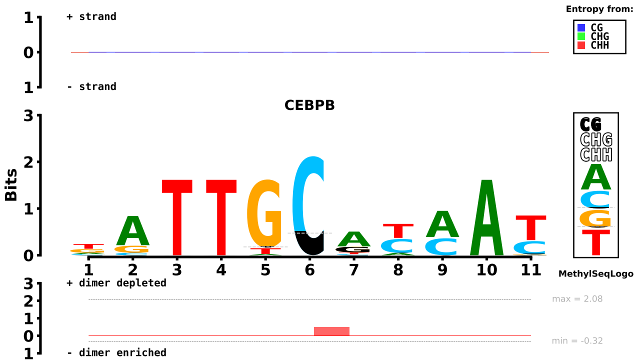


(a) Whole Genome background

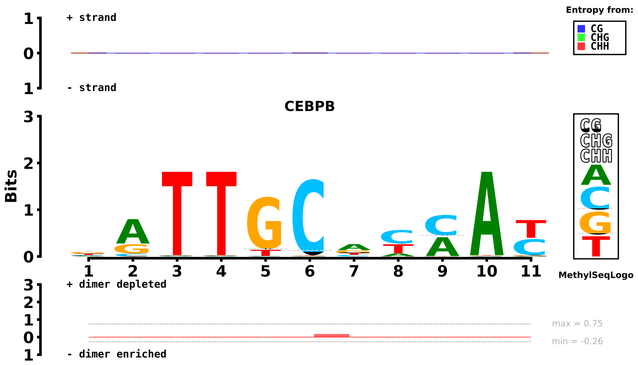


(b) Promoter Region background

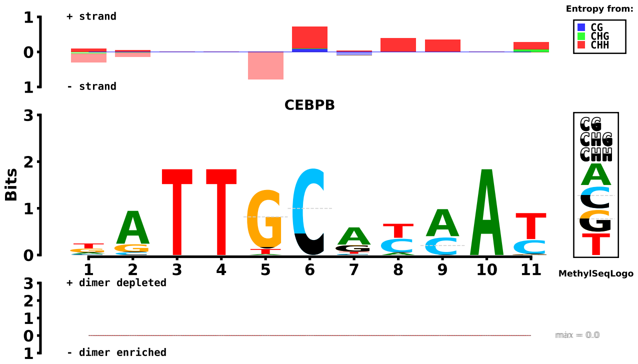


(c) Uniform Background

Figure 5: Transcription factor CEBPB MethylSeqLogos under different background models. (a) Whole-genome (all binding sites against a whole-genome based background), (b) Promoter (promoter region binding sites against a promoter region based background) and (c) Uniform (all binding sites against a uniform background). Data from H1-hESC.

#### Related Visualization Tools

MeDReaders (Wang *et al.*, 2018) is a database summarizing DNA methylation level and TFBS coordinates. MethMotif (Xuan Lin *et al.*, 2018) is a database organizing tissue-specific data. Both of these resources provide methylation aware sequence logos for the convenience of their users. Figure 6 shows MethylSeqLogo compared to the MeDReaders and MethMotif logos.

The MeDReaders sequence logo simply uses `E` to represent methylated cytosine and `C` to represent unmethylated cytosine. This method of representation treats methylated and unmethylated cytosines as being just as different as, for example, guanine and thymidine. This has the unfortunate effect of understating the information in cytosine dominant positions. For example, a position with cytosine in 100% of the binding sites (but methylated 50%) would appear to have the same amount of information as a position with a 50%-50% mix of adenine and thymidine. MeDReaders displays two separate logos; one for highly methylated sites and one for less methylated sites.

MethMotif provides methylation sequence logos split into two images, one for each strand, with bars indicating `CpG` methylation. MethMotif displays `CpG` methylation categorized as methylated ($> 90\%$), heterogeneously methylated (10%–90%), or unmethylated ($< 10\%$); in stacked bars above a traditional sequence logo (Xuan Lin *et al.*, 2018). MethylSeqLogo however, calculates relative entropy from sequence as well as methylation, and forms an integrated logo with `CG` and non-`CG` methylation included.

Finally, we mention a related, but less directly comparable visualization method. Meth-eLogo (Zuo *et al.*, 2014) extends affinity (energy) sequence logos used to visualize energy PWMs (Foat *et al.*, 2006; Stormo, 2013) including methylation contribution as measured by Methyl-Spec-seq experiments (Zuo *et al.*, 2014). Two additional letters (`M` & `W`) are introduced to represent the energetic contribution of adding a 5′-methyl group to a cytosine on the forward or reverse strand respectively. Due to the nature of the affinity data for which it was designed, Meth-eLogo does not show statistical information such as the frequency of methylation in TFBSs or background models.

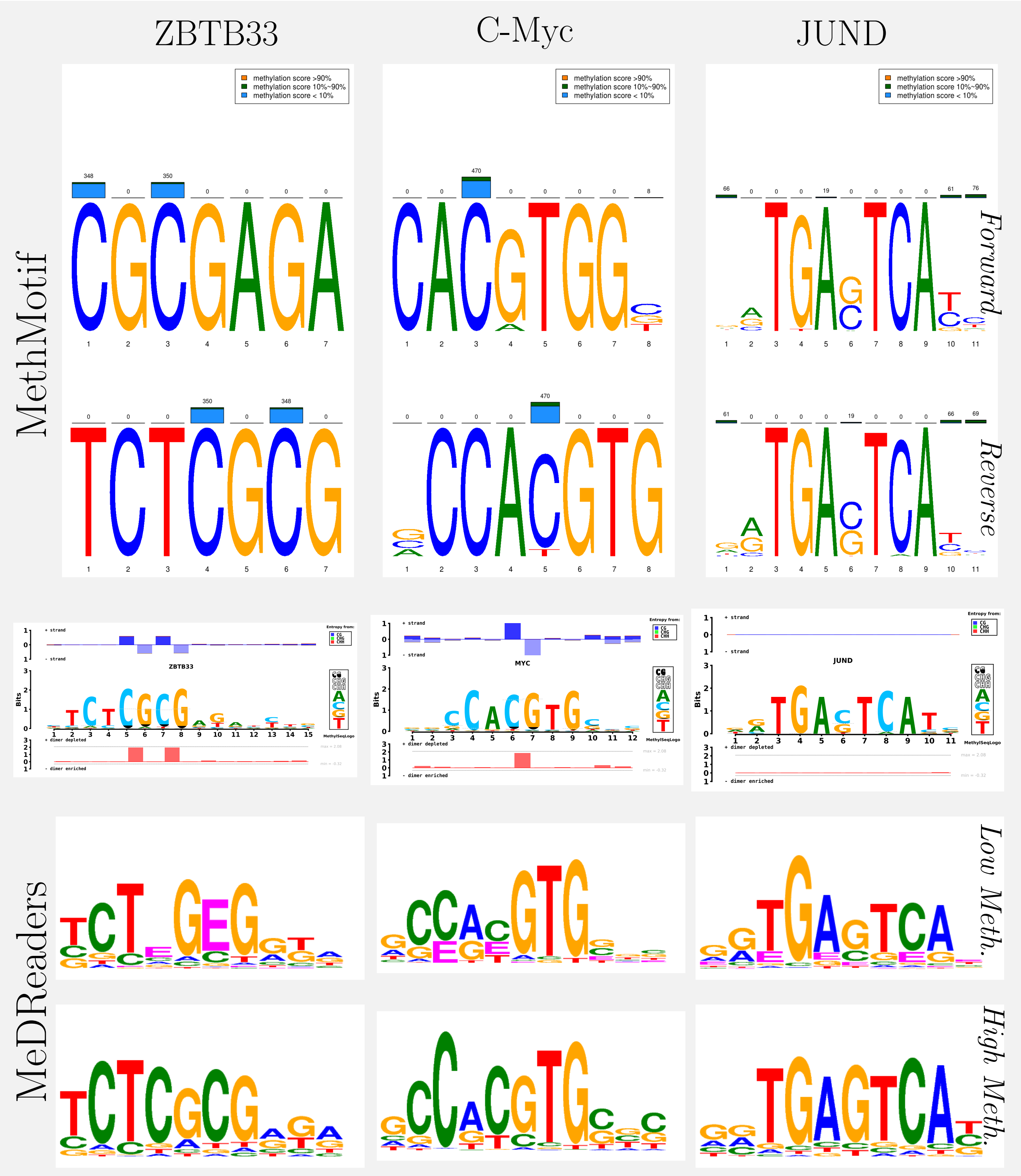


Figure 6: Methylation aware sequence logo methods compared. MethylSeqLogo (middle), MethMotif (top), and MeDReaders (bottom) logos are compared for ZBTB33 (left column), C-Myc (middle column) and JUND (right column). All logos are based on whole genome; HepG2 data for ZBTB33 and JUND, and H1-hESC data for C-Myc.
